# Annual life-history strategy hitchhikes low-light adaptation in a clonal seagrass

**DOI:** 10.64898/2026.06.29.735426

**Authors:** Xiaomei Zhang, Fei Zhang, Zhaxi Suonan, Yu Zhang, Yu-Long Li, Xiaoyu Li, Seung Hyeon Kim, Yi Zhou, Kun-Seop Lee, Lei Yu

## Abstract

While life-history strategies are typically fixed within species, evolutionary transitions between perenniality and annuality can occur. In clonal seagrasses, annual and perennial plants often coexist in the same population, providing a unique model for studying the genetic basis of this transition. Two seagrass *Zostera marina* populations in South Korea display a striking dichotomy: shallow-water sub-populations follow a typical perennial strategy, whereas their deep-water counterparts are annual. Here we show that this shift from perenniality to annuality, potentially caused by the *SAPK7* gene, is genetically coupled with the *CAO* gene, which is under strong positive selection for low-light adaptation. The up-regulation of the *SAPK7* gene triggers early flowering in seedlings, before the formation of any lateral shoots via asexual reproduction. In this special case where the genet contains only one ramet, the post-reproductive death of the ramet is equivalent to the death of the whole genet, which explains the annual phenotype. Our findings reveal a mechanistic example where annuality overcomes perenniality by hitchhiking on a positively selected gene. Given that the ancestral state of plants is perennial, this coupling of annuality with beneficial alleles may represent one of the pathways for the repeated evolution of annual life histories across flowering plants.

---

Plants exhibit a diverse array of life-history strategies^1^. Annual plants complete their life cycle within a single year, with most shoot apical meristems (SAMs) transitioning to floral meristems during the reproductive season, after which the entire plant undergoes senescence and dies before the next growing season. In contrast, biennials and perennials persist for more than one year. This becomes more complex in clonal plants, where a single genetic individual (genet) may consist of numerous physiologically distinct ramets produced via vegetative propagation^2^. In clonal trees, each ramet retains a complete architectural structure, with SAMs organized hierarchically^3,4^. Seagrasses offer a simplified system in which each ramet, or shoot, contains only a single SAM^5^.

Following seed germination, rhizome branching generates new SAMs that develop into independent shoots. During flowering, these SAMs differentiate into floral meristems, leading to shoot death. Theoretically, a genet is perennial as long as at least one ramet remains vegetative. In this study, we introduce a definition of genetic annuals specific to seagrasses: genets in which all ramets flower within the first year following seed germination.

Although life-history strategies are typically fixed at the species level^1^, seagrasses present a notable exception. Some seagrass genets have persisted for centuries or even millennia^6–9^, clearly falling within the perennial category. Conversely, certain seagrass meadows vanish entirely after flowering^10–12^, suggesting an annual strategy. Identifying genetic annuals, however, is technically challenging, as ramet death can result from environmental stress rather than from the vegetative-to-floral transition of SAMs. To investigate the genetic underpinnings of perenniality–annuality transitions in seagrasses, we focus on two populations of the seagrass *Zostera marina* in South Korea (Supplementary Table 1). In each population, deep-water sub-populations completely disappear shortly after flowering, with nearly all flowering individuals consisting of only one single shoot, which meets our definition of genetic annuals. In contrast, adjacent shallow-water sub-populations are perennial.

Chromosomal inversions have been increasingly recognized as key drivers linking suites of phenotypic traits^13–15^. In inversion heterozygotes, recombination is strongly suppressed within the inverted region during meiosis, allowing the entire block of genes to behave as a single “supergene”^16,17^. Such inversions can facilitate local adaptation in the face of gene flow by maintaining favorable combinations of alleles, i.e., adaptive haplotypes, as inherited units. Moreover, because large inversions can lock together multiple genes, they may couple phenotypes that appear otherwise unrelated. In the two Korean *Z. marina* populations, the annual life-history strategy appeared to be tightly linked with adaptation to low-light conditions^10,11^. Here, we tested whether this linkage was mediated by chromosomal inversion.

## Results

### Global *Z. marina* phylogeny

To place the *Z. marina* from East Aisa into the global phylogeny, we conducted whole-genome resequencing for samples from the South Korea and China (Fig. 1a, Supplementary Table 1), with a minimum coverage of 34.83x (Supplementary Data 1). The data were analyzed together with the global *Z. marina* dataset published in Yu et al. (2023)^18^. To be comparable with Yu et al. (2023)^18^, we used the chromosome-level Finnish reference genome^19^. After SNP calling and filtering, a total of 6,525,347 SNPs across 356 samples were obtained.

**Fig 1:**
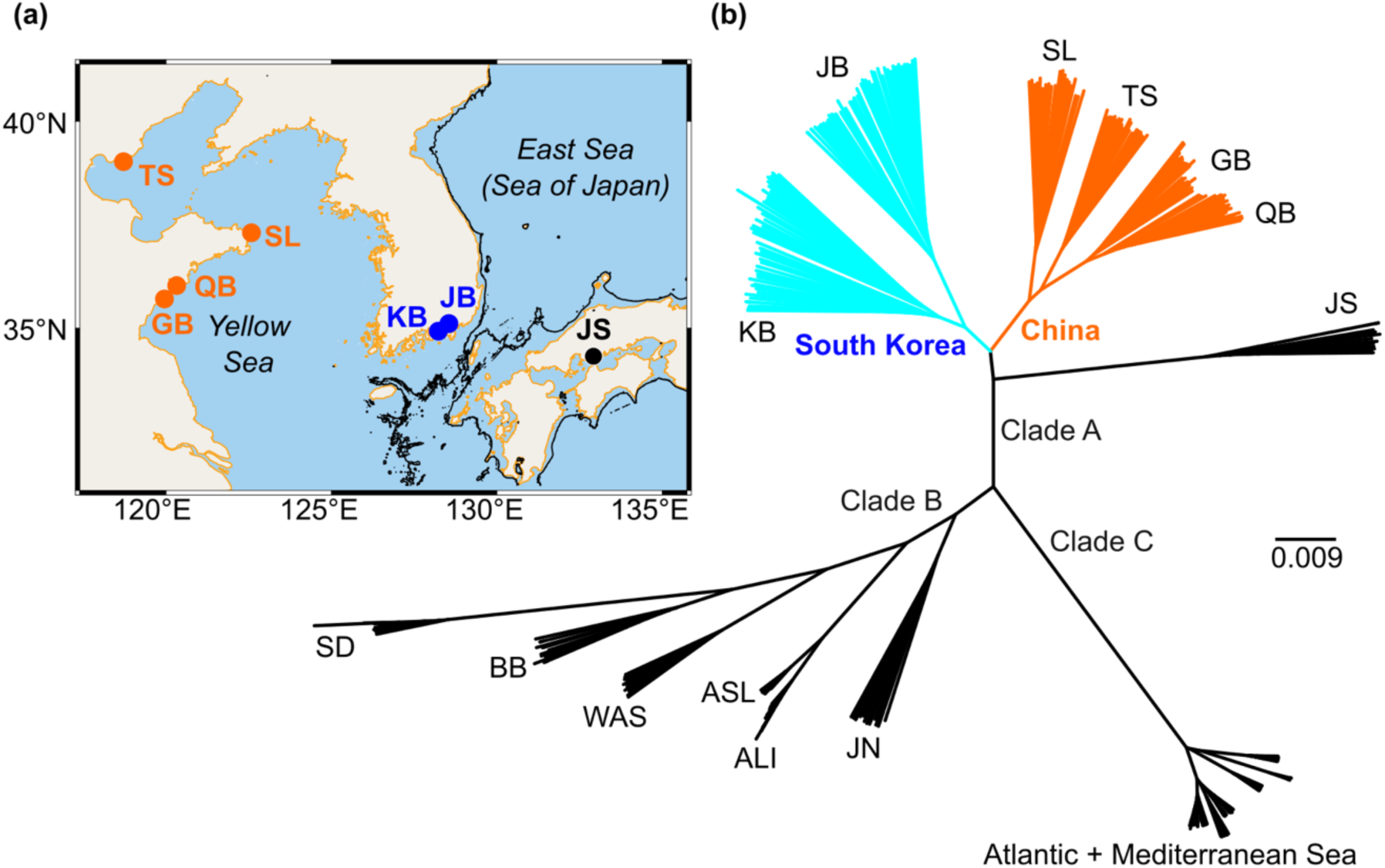
Global phylogeny of the seagrass *Zostera marina*. **a,** The population JS in southern Japan represents the origin of the species. Samples from four populations of China (orange) and two populations of South Korea (blue) are collected and sequenced in this study. A haplotype-resolved genome is constructed for a sample from the population QB. The black lines indicate the −120 m isobath, which roughly represents the coastline during the Last Glacial Maximum (LGM) when the sea level dropped > 120 m. **b,** The sequencing data in this study (blue and orange) are analyzed together with the global sequencing dataset (black) published in Yu et al. (2023)^18^. A Neighbor-Joining (NJ) tree is constructed based on pairwise genetic distance (1 - ibs), which reveals three genetic clades.

In the Neighbor-Joining tree, the samples clustered into three major genetic clades (Fig. 1b). The 4 Chinese populations and the 2 Korean populations clustered together with the population JS from southern Japan, forming the Clade A. The Clade B and C were consistent with the results of Yu et al. (2023)^18^. The Clade A was genetically divergent from the Clade B, the other Pacific clade containing another Japanese population JN from Hokkaido. During the Last Glacial Maximum, the sea level decreased >120 m^20^, and locations of the Chinese or Korean populations in this study were exposed to the air due to the sea-level drop (Fig. 1a). Those populations were likely to be derived from the glacial regium represented by the population JS.

### Chromosomal inversion

To test whether chromosomal inversion was related to the shallow-deep divergence in the two Korean populations, we conducted local PCA (Principal Component Analysis)^21^ for the 86 Korean samples. MDS (Multidimensional Scaling) applied to local PCA results showed how local principal components vary across different regions of the chromosomes (Fig. 2a). The first MDS showed an inversion on Chr02 (1 bp - 7,076,591 bp). Based on the 49,150 SNPs in this region, the first PC explained 62.32% of the variance, which divided the samples into three groups corresponding to genotypes (Fig. 2b). Therefore, the first PC represented the genetic divergence caused by the inversion. We named the homozygous genotypes as INV.00 and INV.11, respectively, and the heterozygous genotype as INV.HET. Correspondingly, the two different states of the inversion site were named as INV.0 and INV.1, respectively. The second PC explained 7.67% of the variance, separating the two different geographic locations (Fig. 2b).

**Fig 2:**
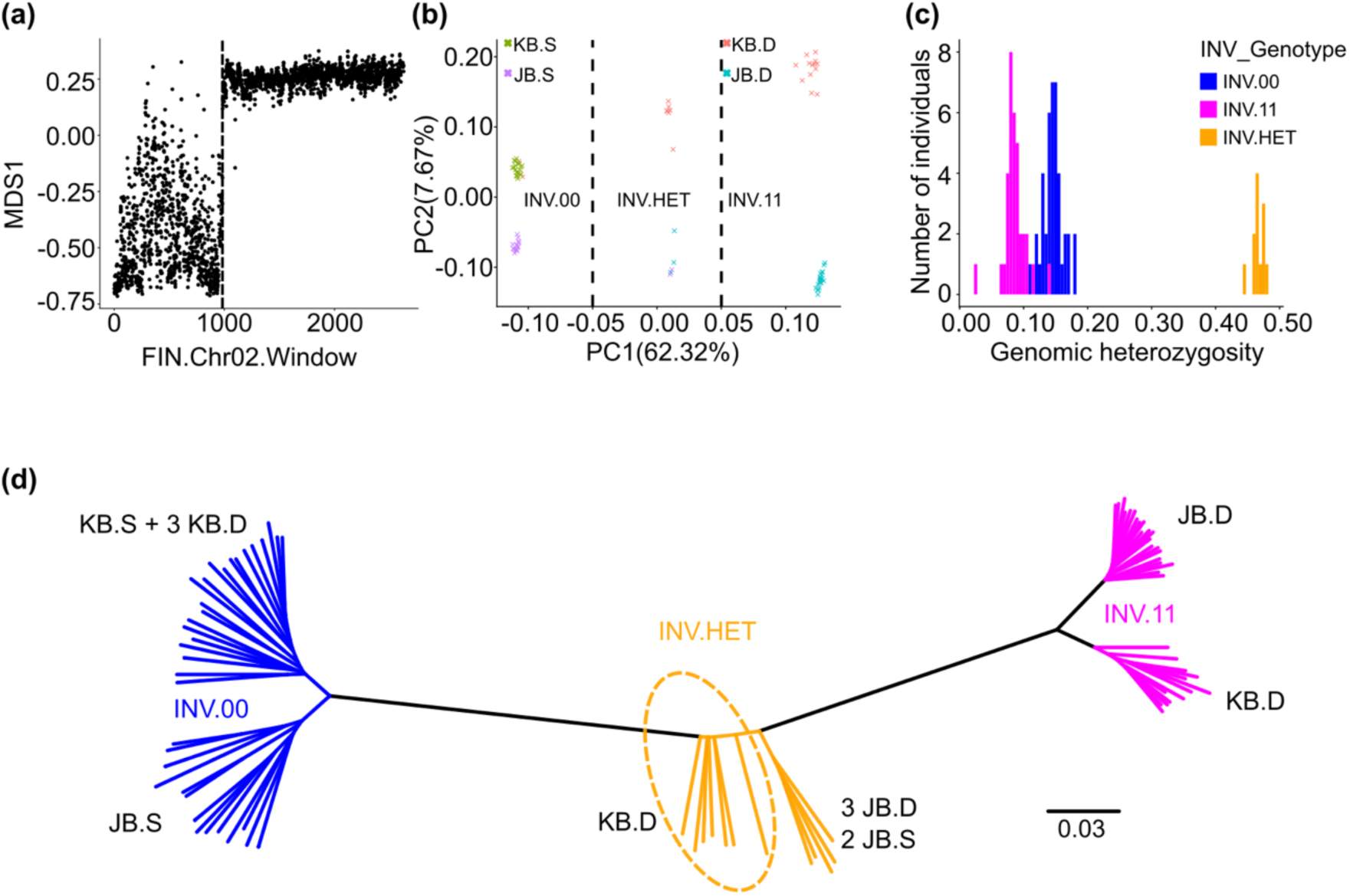
A 7-Mb chromosomal inversion underlying the shallow-deep divergence in the South Korea. **a,** MDS1 shows how local principal components vary across different regions of the Chr02 in the Finnish genome^19^. The vertical dashed line is placed at 7,076,591 bp. The region 1-7,076,591 bp represents a chromosomal inversion. **b,** Based on the SNPs located in the inversion region, the first PC divides the Korean samples into three groups, corresponding to different genotypes. KB.S and JB.S represent the shallow-water sub-populations, while KB.D and JB.D represent the deep-water sub-populations. **c,** Genetic diversity of the three genotypes are assessed by genomic heterozygosity. The inversion heterozygotes show the highest genomic heterozygosity. As for the two homozygous genotypes, the genomic heterozygosity of the genotype INV.00 is higher, indicating that the inversion state INV.0 is the ancestral state. **d,** A Neighbor-Joining tree is constructed based on pairwise genetic distances (1 - ibs). The genotype INV.11 is only distributed in deep-water regions.

To figure out whether INV.0 or INV.1 represented the ancestral state, we assessed the genetic diversity of the three genotypes. INV.HET was expected to show the highest genetic diversity, as it contained polymorphisms from both states. As for the two homozygous genotypes, the homozygote for the ancestral state was supposed to show higher genetic diversity. In our case, the genotype INV.00 showed higher genomic heterozygosity than the genotype INV.11 (Fig. 2c), indicating that INV.0 represented the ancestral state. The Neighbor-Joining tree was consistent with the PCA results (Fig. 2d). The genotype INV.00 was distributed mostly in the shallow-water sites (KB.S and JB.S), except for the 3 samples from the deep-water site of the population KB (i.e., KB.D). In contrast, the genotype INV.11 was only found in deep-water sites (KB.D and JB.D). As for the heterozygote INV.HET, most of the samples were distributed in the deep-water sites (KB.D and JB.D), except for the two samples from the shallow-water site of the population JB (i.e., JB.S).

### Global distribution of the chromosomal inversion

To demonstrate the global distribution of the chromosomal inversion, we constructed the global *Z. marina* phylogeny based on the 256,217 SNPs in the inversion region (FIN.Chr02: 1 bp - 7,076,591 bp). The different genotypes of the inversion formed three clusters, and the cluster representing the genotype INV.HET was identified by their higher genetic diversity (Supplementary Fig. 1). The inversion state INV.1 was only present in the Chinese population QB, the two Korean populations KB and JB, and the population JS in southern Japan (Fig. 3). Apart from those 4 populations, all the rest populations were fixed with the state INV.0, which also supported that INV.0 was the ancestral state. The topology within the genotype INV.00 was consistent with that in the global phylogeny (Fig. 1b).

**Fig 3:**
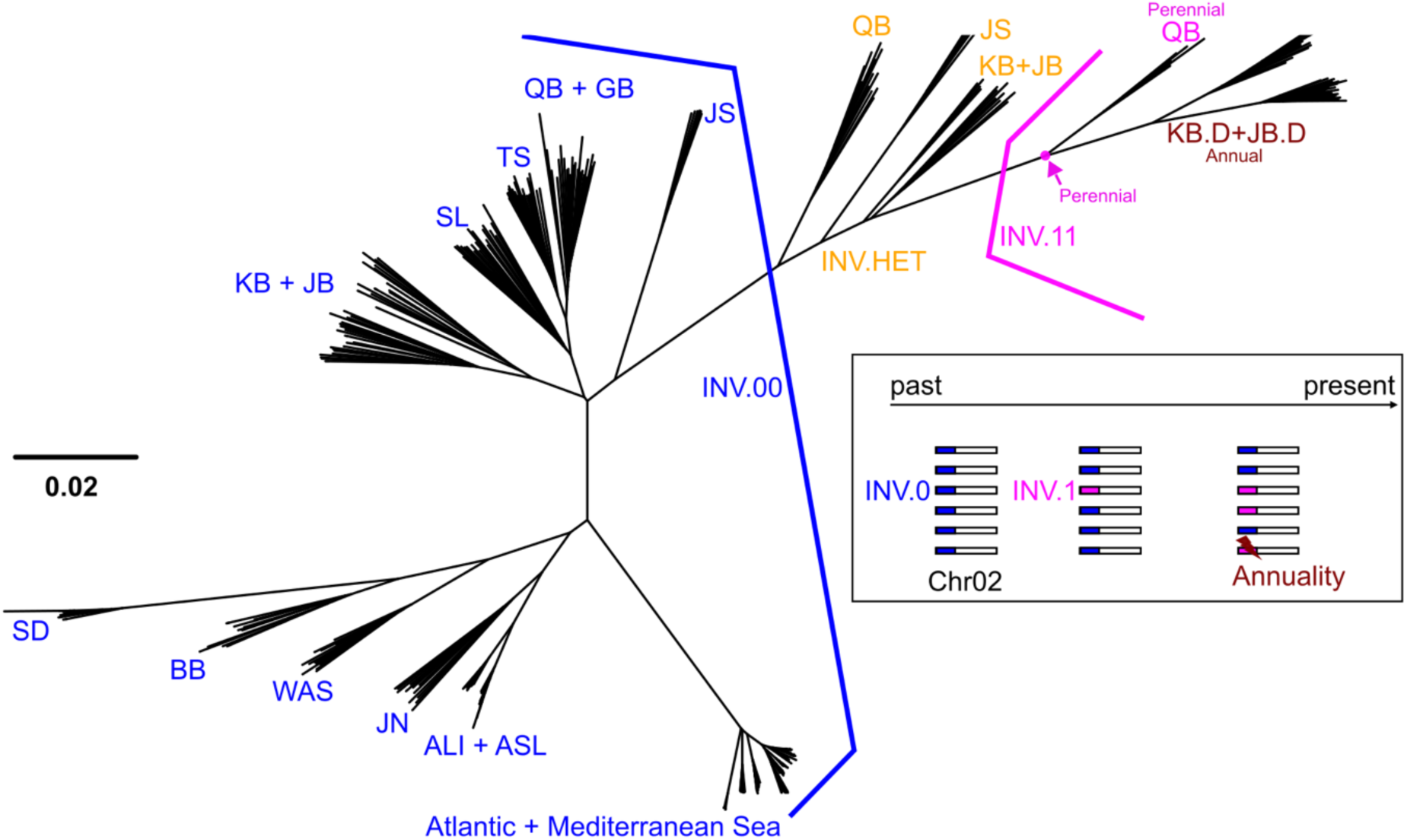
Global distribution of the 7-Mb chromosomal inversion. A Neighbor-Joining tree is constructed based on the SNPs located in the inversion region (FIN.Chr02: 1-7,076,591 bp). The inversion heterozygotes are identified based on their high genomic heterozygosity (Supplementary Fig. 1). The genotype INV.11 contains both annual and perennial samples, indicating that annuality emerged after the inversion event.

We further confirmed the existence of the inversion by assembling the haplotype-resolved genomes of an INV.HET sample from the population QB (Fig. 4). The chromosome numbering in the QB genome was based on the length, therefore it was not surprising that the numbering did not match the Finnish genome. The Chr02 in the Finnish genome corresponded to the Chr05 in the QB genome. For the coordinates 1 bp – 6 Mb in Chr02 of the Finnish genome, the two haplotypes of the QB genome showed different inversion states (Fig. 4). The detected break point of the inversion here (i.e., 6 Mb) was slightly different from that in the Korean populations (i.e., 7,076,591 bp). A 1-Mb difference was not likely to be caused by technical differences (SNP-based vs. assembly-based methods). Therefore, it might be due to their divergent evolutionary histories (Fig. 1b). To obtain the coordinates of the inversion in the QB genome, we mapped the data of the Korean samples against the first haplotype of the QB genome, i.e., QB.hap01. After SNP calling and filtering, a total of 1,116,735 SNPs were kept. Using local PCA, the coordinate of the inversion was found to be Chr05: 28,209,630 bp – 38,049,035 bp (Supplementary Fig. 2).

**Fig 4:**
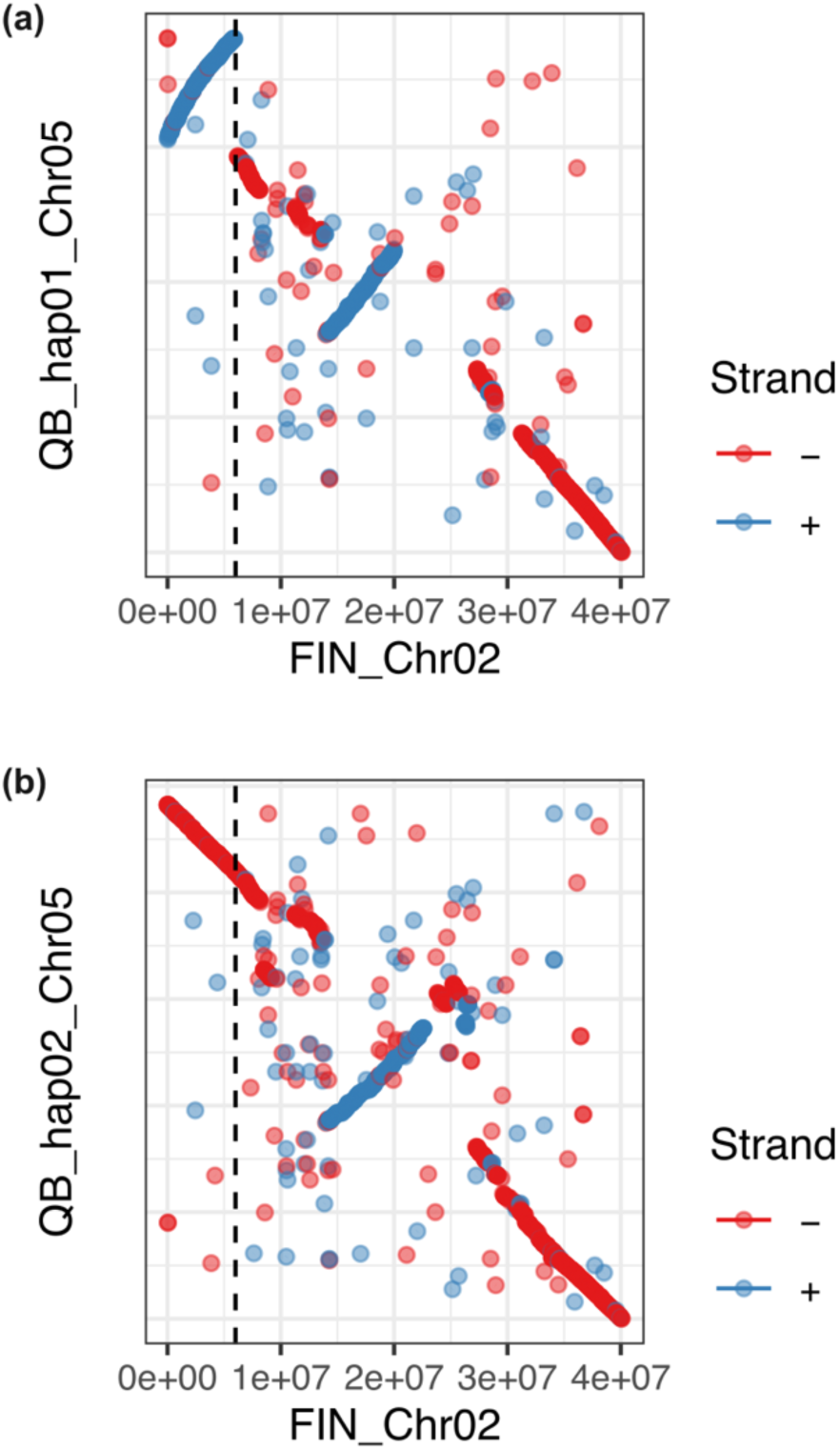
DNA sequence alignment. The x axis represents the Chr02 of the Finnish genome^19^, and the y axis represents the haplotype no. 1 (**a**) and no. 2 (**b**) for Chr05 of the genome assembled in this study for a sample collected from the population QB. All the maximal unique matches of 20 bp between the two input sequences are identified, which are further clustered into closely grouped sets. All clusters with a length of > 1000 bp are shown without any filtering. The vertical dashed line is placed at 6 Mb. The QB sample is an inversion heterozygote for the region Chr02:1 bp – 6 Mb in the Finnish genome.

### Genetic basis for low-light adaptation

All the samples with the homozygous genotype INV.11 in the two Korean populations were distributed in the deep water (4.5 m, Supplementary Table 1), indicating that the state INV.1 might adapt to the low-light environment. To detect signatures of positive selection, we calculated Tajima’ D for the INV.11 samples^22^. We first checked whether there were ramets belonging to the same clone (Supplementary Table 2), as it would bias the Tajima’ D analysis, and we found that all 34 INV.11 samples represented unique genets. The inversion region was divided into 10,000-bp bins, and one bin (FIN.Chr02: 4140000 bp – 4150000 bp) showed Tajima’ D value < −2.5 (D = −2.68, p < 0.05, Supplementary Fig. 3), which contained 3 genes (*Zosma02g04960*, *Zosma02g04970*, and *Zosma02g04980*). The gene *Zosma02g04960* encoded chlorophyllide *a* oxygenase (CAO). To exclude the potential reference bias on the Tajima’s D calculation, we replaced the Finnish genome with the QB genome (QB.hap01) to conduct the Tajima’s D analysis independently (Fig. 5a). Two adjacent bins showed D values < −2.5, containing 7 genes in total, and one of the genes (*Zmar_015735*) also encoded chlorophyllide a oxygenase (CAO). Since the QB genome was assembled and annotated independently, it was reassuring that the *CAO* gene was identified again.

**Fig 5:**
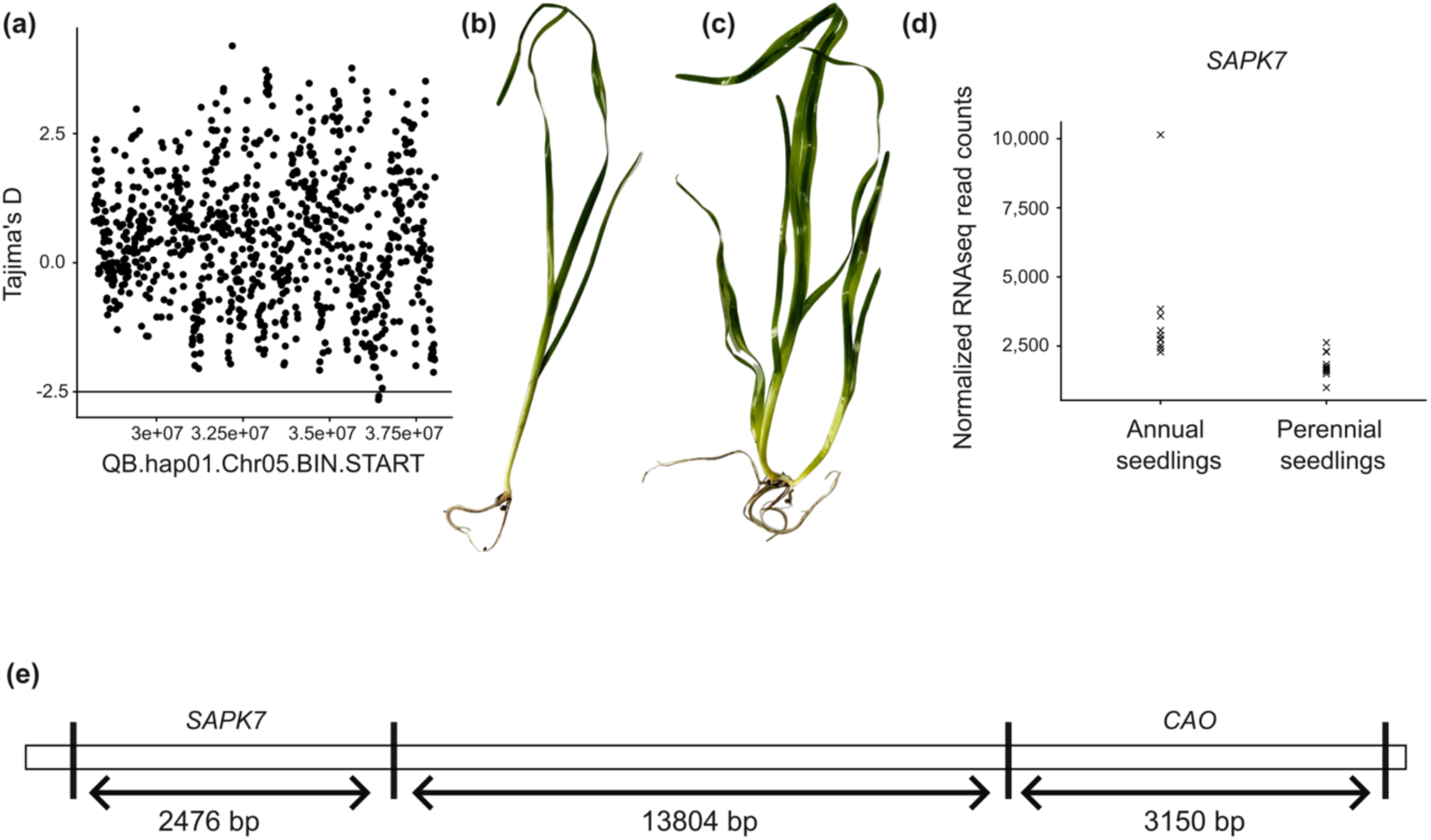
Annual life-history strategy hitchhikes low-light adaptation. **a,** The inversion region (QB.hap01.Chr05: 28,209,630 – 38,049,035) is divided into 10,000-bp bins for calculating Tajima’ D. Two adjacent bins (Chr05: 36,410,000-36,430,000 bp) have D values < −2.5, indicating positive selection. The two bins contain 7 genes in total, and one of the genes encodes chlorophyllide a oxygenase (CAO), which plays an important role in low-light adaptation of plants. **bcd,** To identify the genes related to the annual life history, we look at the gene expression in the inversion region. Two types of plants are selected. The first group consists of seedlings with a single reproductive shoot at the pre-flower bud stage (i.e., Annual seedlings, **b**), while the second group consists of seedlings that have produced a lateral shoot and both shoots remain vegetative (i.e., Perennial seedlings, **c**). A total of 35 genes are differentially expressed between the two groups (adjusted p < 0.01, Supplementary Data 3), including a *SAPK7* gene (**d**). The up-regulation of the *SAPK7* gene triggers early flowering in seedlings.In a special case where the genet contains only one ramet, the post-reproductive death of the ramet is equivalent to the death of the whole genet, which explains the annual phenotype. **e,** The *CAO* gene and the *SAPK7* gene are separated by only 13,804 bp.

To test whether the significantly negative D value was a unique feature for the Korean samples, we conducted the same analysis for the INV.11 samples from the population QB in China. To increase the number of the INV.11 samples, we sequenced another 80 samples from the population QB. None of the QB samples belonged to the same genet (Supplementary Table 2). The number of samples with the genotypes INV.00, INV.HET, and INV.11 was 28, 35, and 17, respectively. Based on those counts, the Hardy-Weinberg equilibrium could not be rejected, which indicated that the inversion was not under selection in the population QB. We then calculated the Tajima’s D values using the 22 INV.11 samples. The values for the same two bins detected above were 1.65 and 2.00, respectively, showing no signature of positive selection. In conclusion, the significantly negative Tajima’s D value was only observed in the Korean INV.11 samples. There were several scenarios that might lead to significantly low D values, such as purifying selection, positive selection, and population expansion after a bottleneck. In our case, the most plausible explanation would be positive selection.

To identify the SNPs in the *CAO* gene related to the low-light adaptation, we conducted comparison between the INV.11 samples from South Korea and the Chinese population QB. A total of 63 SNPs were found in the *CAO* gene (Supplementary Data 2). The Korean samples were almost fixed with one haplotype, as almost all the samples shared the same homozygous genotype at each SNP (Supplementary Data 2). This indicated positive or purifying selection, consistent with the Tajima’s D analysis. The samples from the population QB showed higher levels of polymorphism. A total of 7 SNPs changed the amino acids, and another 5 SNPs were found in the 5’ or 3’ UTRs. There was also a SNP leading to a stop codon. For the 3-bp locus from 36,420,409 bp to 36,420,411 bp in the Chr05 of the genome QB.hap01, the samples from the population QB showed two different haplotypes TAC and TAA, with 14 out of 22 samples fixed with TAA, one of the stop codons. In contrast, almost all the Korean samples (32 out of 34 samples) were fixed with the alternative haplotype TAC.

The *CAO* gene plays an important role in low-light adaptation of plants by optimizing the light-harvesting capacity. The enzyme chlorophyllide a oxygenase (CAO) converts chlorophyll a to chlorophyll b^23^, which is a key structural component of the light-harvesting complexes (LHCs), the “antennae” of the photosystems. An increased supply of chlorophyll b allows the plant to enlarge the antennae of the LHC proteins, so that more photons will be captured.

### Potential genetic basis for annuality

The global distribution of the chromosomal inversion rejected our hypothesis that the annual life-history strategy and low-light adaptation were linked by the chromosomal inversion (Fig. 3), as there were INV.11 samples (from the population QB) that were perennial. The samples from the population QB were collected after the flowering season (Supplementary Data 1). Since annual plants were supposed to die during the first flowering season, the INV.11 samples from the population QB were expected to be perennial. Hence, the inversion state INV.1 did not directly correspond to annuality, and the annual life-history strategy emerged after the born of the inversion state INV.1 (Fig. 3).

To identify the genes potentially related to the annual life history, we furthered looked at the gene expression in the inversion region, i.e., QB.hap01.Chr05: 28,209,630 – 38,049,035. We conducted RNA sequencing for two groups of samples. The first group consisted of seedlings with a single reproductive shoot at the pre-flower bud stage (i.e., Annual seedlings, Fig. 5b)^24^, while the second group consisted of seedlings that had produced a lateral shoot and both shoots remained vegetative (i.e., Perennial seedlings, Fig. 5c). A total of 35 genes in the inversion region were differentially expressed between the two groups (p < 0.01, Supplementary Data 3).

One gene was *Zmar_015731*, which encoded Serine/threonine-protein kinase SAPK7. Liu et al. (2024)^25^ systematically analyzed all ten members of the *SAPK* family in rice. By creating mutations in each gene with gene-editing technology, they found that mutants with a non-functional *SAPK7* gene showed a delayed flowering phenotype. Therefore, the higher expression of the *SAPK7* gene might lead to early flowering. The *SAPK7* gene was up-regulated in the “Annual seedlings” compared with the “Perennial seedlings” (p< 0. 01, Fig. 5d). It was likely that the annual life-history strategy was caused by early flowering. Under a scenario where the seedling started flowering before the formation of any lateral shoots through asexual reproduction, the only SAM differentiating into floral meristem means the death of the plant. In addition, the *SAPK7* gene was only 13,804 bp away from the *CAO* gene (Fig. 5e), which explained why the annuality and low-light adaptation were linked together.

## Discussion

Our understanding of the evolutionary and molecular transitions between perenniality and annuality in plants remains limited. In Brassicaceae, the shift from polycarpic perennial to biennial and annual flowering is governed by the dosage of three closely related MADS-box genes^26^. In rice, Dai et al. (2026)^27^ identified the *Endless Branches and Tillers (EBT1)* locus, which drives floral reversion and vegetative propagation, thereby underpinning perennial growth in wild relatives. Here, we propose that early flowering potentially triggered by the *SAPK7* gene, may constitute an independent mechanism for conferring annuality in seagrasses. Specifically, flowering occurs before the establishment of lateral shoots, leading to whole-plant senescence and thus completing the annual life cycle.

Given that annuality and perenniality are determined by one single gene, in a mixed population with both annuals and perennials, the ratio of the annual to perennial individuals should theoretically remain stable in the seed-derived cohorts; however, perennials possess the additional advantage of overwintering and contributing to the gene pool of the subsequent generations. This would predict a progressive increase of the perenniality allele frequency over time (Supplementary Fig. 4). Contrary to this expectation, we find that the annual strategy dominates deep-water sub-populations of *Zostera marina* in two Korean populations. This counterintuitive pattern is explained by the annuality hitchhiking on the *CAO* gene, which is under positive selection in low-light environments (Fig. 5a). The physical coupling effectively links the annual phenotype with an adaptive advantage under light limitation, allowing annuals to overcome perennials.

Annuality in *Z. marina* has been observed across a broad spectrum of stressful environments, where populations are subject to ice scour, grazing, aerial exposure, thermal stress, hypoxia/anoxia, low-light conditions, or combinations thereof^10,28–33^. From an ecological standpoint, the annual strategy may enhance meadow persistence, resilience, and recovery in disturbed settings by promoting seed-based regeneration, rapid population turnover, and effective recolonization^11,12,34^. Our study offers an evolutionary example in which linkage to positively selected genes provides a mechanism for annuality to overcome the competitive advantage typically associated with perenniality.

The genomic coupling of the *CAO* gene and the *SAPK7* gene produces annual plants pre-adapted to low-light conditions, enabling them to occupy a “refuge” niche where light is limiting. Moreover, these plants complete germination, flowering, and seed set within a few months, allowing for rapid fitness feedback—a cycle that operates on a seasonal rather than multi-year timescale. This genetic package not only sheds light on the evolutionary persistence of annuality in clonal marine angiosperms, but also represents a valuable resource for future restoration efforts in *Z. marina* meadows^35^.

## Methods

### Sample collection and whole-genome resequencing

Samples from the two Korean populations were collected by SCUBA. As for the four Chinese populations, samples were collected at low tide when the water is less than knee-deep. Bulk DNA from the meristematic region and the basal portions of the leaves, was extracted using a CTAB method. DNA quality was checked following the requirements of Biomarker Technologies (BMK). Libraries with insert size around 350 bp were prepared, which were then sequenced with DNBSEQ-T7 (PE150).

### Assembling and annotating the genome for a sample from the population QB

A single genet with multiple shoots connected by the rhizome was collected from Qingdao Bay (QB), China (36°3′N, 120°19′E). DNA was extracted with a CTAB method, which was used for Pacbio HiFi sequencing (15 kb library, Pacbio Sequel II system) and standard whole-genome resequencing (350-bp insert size, PE150, DNBSEQ-T7). Hi-C library was also constructed, which was sequenced on DNBSEQ-T7 (PE150). Different types of tissues were used for RNA sequencing. The extracted RNA was used as template to generate the first strand of cDNA, which was then used as template to generate the second strand. The libraries were sequenced on Illumina Novaseq 6000 platform (PE150). In addition, the RNA sequencing was also conducted with Pacbio Sequel II system.

PacBio HiFi reads of *Zostera marina* were assembled into contigs using HIFIASM V0.19.8^36^ with the parameters -s 0.5 --ul-cut 20000 --ul-rate 0.02. Haplotypic duplications were subsequently purged using PURGE HAPLOTIGS V1.1.2^37^. The purged contigs were then scaffolded using Hi-C reads with YAHS V1.2.2^38^. The resulting scaffolds were manually curated and corrected using JUICEBOX V1.11.08 (https://github.com/aidenlab/Juicebox). Assembly quality was assessed by mapping short reads back to the assembly using BWA V0.7.17^39^, and genome completeness was evaluated using BUSCO V6.0.0^40^ against the embryophyta_odb12 dataset.

Gene prediction was performed by integrating evidence from *ab initio* predictors, protein homology, and transcript alignments. First, a *de novo* repeat library was constructed using EDTA V2.2.0^41^, and repeat sequences (excluding low-complexity regions) were soft-masked. Second, BRAKER3 V3.0.6^42^ was utilized to train the gene prediction tools GENEMARK-ETP V1.00^43^ and AUGUSTUS V3.4.0^44^, and to generate *ab initio* predictions based on RNA-seq data and Viridiplantae protein homology data from ORTHODB V10.

Third, PacBio Iso-Seq reads were processed and clustered into full-length transcripts using the ISO-SEQ V4.0.0 pipeline (https://github.com/PacificBiosciences/IsoSeq). Short RNA-seq reads were *de novo* assembled using TRINITY V2.15.1^45^. The combined transcriptome assemblies were then passed to PASAPIPELINE V2.5.3^46^ to obtain high-quality gene structures. Fourth, homologous evidence from the aforementioned ORTHODB protein sequences and RNA-seq reads was extracted by aligning them to the genome using METAEUK^47^, MINIMAP2 V2.24^48^, and STRINGTIE V2.2.1^49^. Finally, the *ab initio* predictions, protein alignments, and transcript alignments were integrated into weighted consensus gene models using EVIDENCEMODELER V2.0.0^50^, as implemented in FUNANNOTATE V1.8.15 (https://github.com/nextgenusfs/funannotate). Subsequently, PASAPIPELINE was utilized to update the untranslated region (UTR) annotations based on the above combined transcriptome assemblies.

NUCMER V4.0.0rc1^51^ was used to align the two haplotype-level genomes of the QB sample to the Finnish genome (--mum -c 1000)^19^. A custom R script was used to visualize the alignment results (https://github.com/leiyu37/Hitchhiking_Zm/tree/main/07_nucmer_hap01Fin). The Chr05 of the QB genomes corresponded to the Chr02 of the Finnish genome. At around 6 Mb of the FIN.Chr02, QB.Chr05.hap01 showed a break point for an inversion, while QB.Chr05.hap02 not (Fig. 4). Therefore, the QB sample was an inversion heterozygote for the region FIN.Chr02: 1bp – 6 Mb.

### SNP calling and filtering

Either the Finnish genome^19^ or the QB.hap01 (assembled in this study) was used as the reference genome. The genomes were indexed with BWA V0.7.19 (index)^39^, SAMTOOLS V1.13 (faidx)^52^, and GATK V4.6.2.0 (CreateSequenceDictionary)^53^. The quality of the raw reads was assessed by FastQC (https://www.bioinformatics.babraham.ac.uk/projects/fastqc/). BBDUK (https://jgi.doe.gov/data-and-tools/bbtools/bb-tools-user-guide/bbduk-guide/) was used to remove adapters and for quality filtering (https://github.com/leiyu37/Hitchhiking_Zm/tree/main/01_bbduk), discarding sequence reads (1) shorter than 50 bp after trimming (minlength = 50), and (2) with average quality <20 after trimming (maq = 20). FastQC was used for second round of quality check for the clean reads. Sequencing coverage was calculated for each sample (Supplementary Data 1).

A Genomic Variant Call Format (GVCF) file was generated for each sample (https://github.com/leiyu37/Hitchhiking_Zm/tree/main/02_fq2hap). The quality-filtered reads were mapped against the reference genome using BWA V0.7.19 (MEM), and then alignments were converted to BAM format and sorted using SAMTOOLS V1.13. The MarkDuplicates module in GATK V4.6.2.0 was used to identify and remove duplicate reads in the BAM files. HaplotypeCaller (GATK4) was used to generate a GVCF file for each sample. All the GVCF files were combined by CombineGVCFs (GATK4), and GenotypeGVCFs (GATK4) was used to call genetic variants. Then, a filtering workflow was used to obtain high-quality SNPs (https://github.com/leiyu37/Hitchhiking_Zm/tree/main/04_FINref/50_global/05_filter). VariantsToTable (GATK4) was used to extract INFO annotations. SNPs were filtered by VariantFiltration (GATK4) based on the following parameters: MQ, FS, QD, MQRandSum, ReadPosRandSum, SOR, and DP. Only SNPs were kept, while the other variant types were removed (GATK4 SelectVariants). VCFTOOLS V0.1.16 (Danecek et al. 2011) was used to convert individual genotypes to missing data when GQ < 30 or DP < 10. Only bi-allelic SNPs with at least one minor allele were kept.

### Neighbor-Joining (NJ) tree and clonemate detection

PLINK V1.9.0-b.7.8^54^ was used to calculate pairwise genetic distances (1-ibs), and MEGA V10.2.6^55^ was used to construct a Neighbor-Joining (NJ) tree based on the genetic distance matrix. FIGTREE V1.4.4 (https://github.com/rambaut/figtree.git) was used for visualizing the phylogenetic tree. Clonemate pairs were detected using a method based on shared heterozygosity (https://github.com/leiyu37/Detecting-clonemates)^56^. The SH index was calculated for each pair of samples, and the pairs with SH > 0.95 represented clonemates (Supplementary Table 2).

### Detecting chromosomal inversions

LOCALPCA (https://github.com/petrelharp/local_pca.git)^21^ was used to detect chromosomal inversions based on the MDS (Multidimensional Scaling), and an inversion was identified as the region dividing the samples into three separate clusters. Principal Component Analysis (PCA) and NJ tree were constructed based on the SNPs located in the inversion regions to genotype the samples. The inversion heterozygotes were distributed in between the two homozygous genotypes, and they were identified by the high genomic heterozygosity (https://github.com/leiyu37/Hitchhiking_Zm/tree/main/04_FINref/50_global/07_het).

### Detecting signals of positive selection

Th inversion region was divided into 10,000-bp bins for calculating the Tajima’s D^22^. The calculation was conducted with VCFTOOLS V0.1.16 (--TajimaD 10000). Bins with Tajima’s D smaller than −2.5 was considered under positive selection (https://github.com/leiyu37/Hitchhiking_Zm/tree/main/04_FINref/51_extractKorea/06_t ajimaD). The relevant gene information was extracted from the annotation files (gff3 format). The *CAO* gene showed strong signals of positive selection, and the SNPs located in the CAO gene were annotated with SNPEFF V5.3a (http://pcingola.github.io/SnpEff/).

### RNA sequencing analyses

Samples were collected from the Jindong Bay (JB, Supplementary Data 1) by SCUBA. We focused on two types of samples: the first type consisted of seedlings with a single reproductive shoot at the pre-flower bud stage (Fig. 5b), while the second type consisted of seedlings that had produced a lateral shoot and both shoots remained vegetative (i.e., Fig. 5c). Bulk RNA from the meristematic region and the basal portions of the leaves was used for RNA sequencing. The extracted RNA was used as template to generate the first strand of cDNA, which was then used as template to generate the second strand. The libraries were sequenced on DNBSEQ-T7 (PE150), and the data were analyzed with DESEQ2^57^. Differentially expressed genes between the two types were identified based on the adjusted p values (p < 0.01, https://github.com/leiyu37/Hitchhiking_Zm/tree/main/09_DESeq2).

## Competing interests

None declared.

## Author contributions

Zhou, K-S.L., and L.Y. conceived and designed this study; X.Z. collected and sequenced the samples from China; F.Z., Z.S., and S.H.K collected and sequenced the samples from South Korea; Y. Zhang and Y-L.L. assembled and annotated the genome for a sample from the population QB; X.L. identified the plant types for the RNA sequencing; L.Y. analyzed and interpreted the sequencing data; X.Z. and L.Y. wrote the manuscript. All authors commented on earlier versions of the manuscript. X.Z., F.Z., and S.Z. contributed equally to this work.

## Data availability

The raw sequencing data for the QB genome are deposited in NCBI under the accession no. PRJNA1472446. The resequencing data and the RNAseq data are deposited in NCBI under the accession no. PRJNA1474806. The scripts used in this study are deposited on Github (https://github.com/leiyu37/Hitchhiking_Zm). Genome and annotation for the QB sample are deposited in figshare (https://doi.org/10.6084/m9.figshare.32797473).

## Notes

### Competing Interest Statement

The authors have declared no competing interest.

