## Supplementary material for "Annual life-history strategy hitchhikes low-light adaptation in a clonal seagrass": SupplementaryInformation.pdf

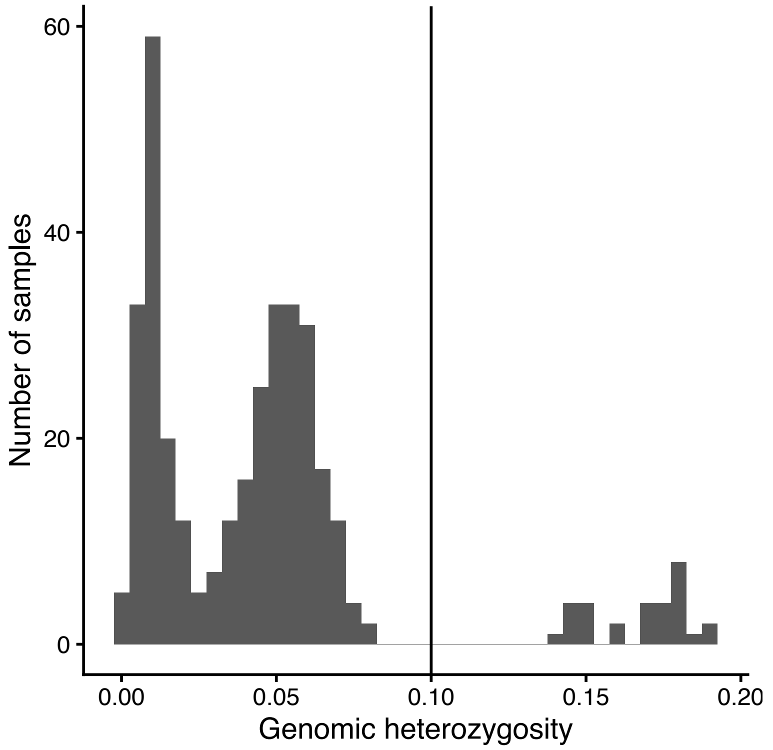


**Supplementary Figure 1: Genomic heterozygosity based on the SNPs located in the inversion region (FIN.Chr02: 1 bp - 7,076,591 bp).** The threshold of 0.1 is used to distinguish between homozygous and heterozygous genotypes. Samples with genomic heterozygosity > 0.1 are inversion heterozygotes.

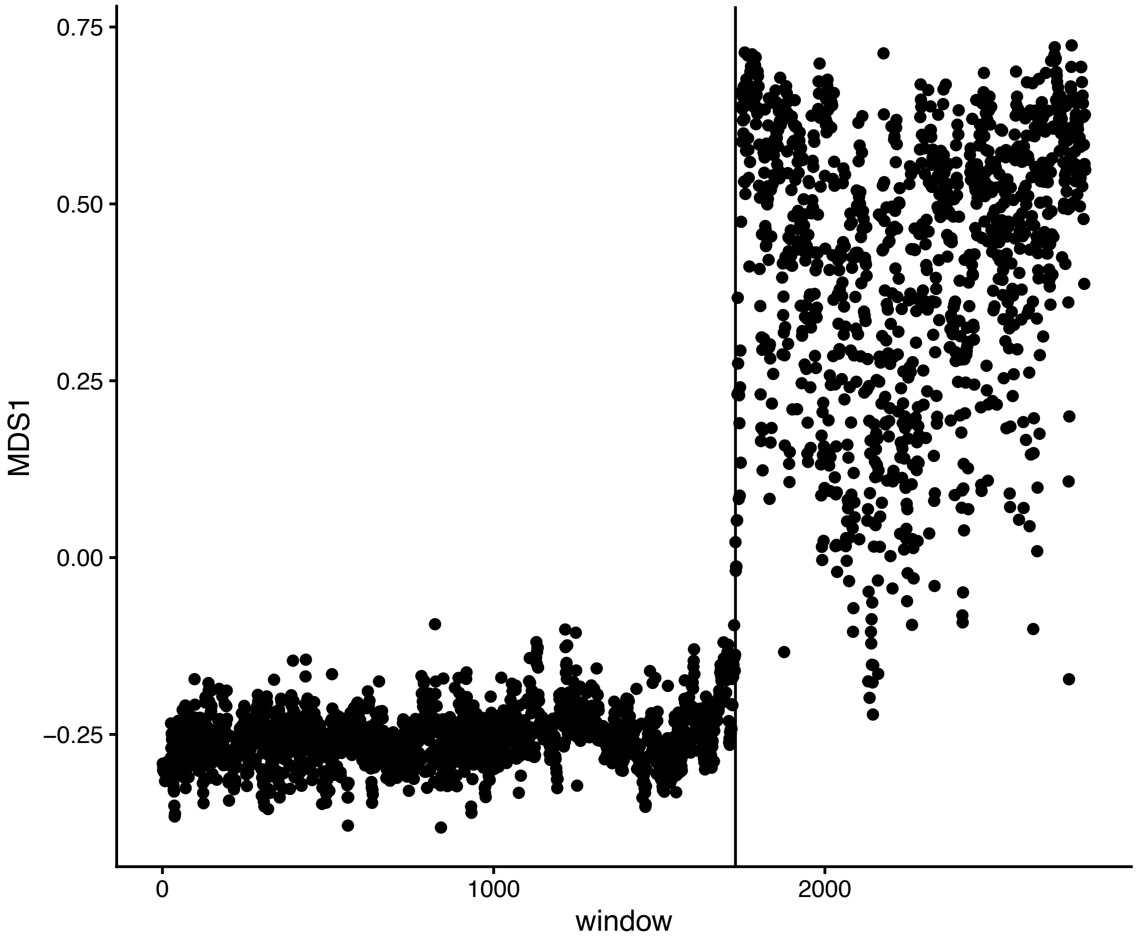


**Supplementary Figure 2: MDS1 along the Chr05 of the genome QD.hap01.** MDS1 shows how local principal components vary across different regions. The vertical dashed line is placed at 28,209,630 bp. The region 28,209,630 – 38,049,035 bp represents a chromosomal inversion.

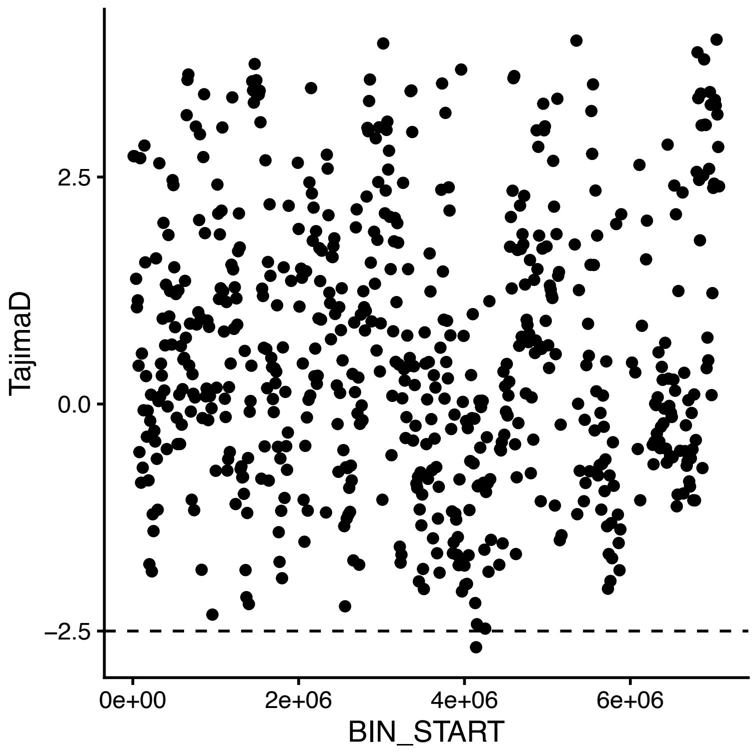


**Supplementary Figure 3: Tajima's D based on the SNPs located in the inversion region (FIN.Chr02: 1 bp - 7,076,591 bp).** The inversion region is divided into 10,000-bp bins for calculating Tajima' D. One bin (4,140,000-4,150,000 bp) has D values < -2.5, indicating positive selection. The bin contains 3 genes in total, and one of the genes encodes chlorophyllide a oxygenase (CAO), which plays an important role in low-light adaptation of plants.

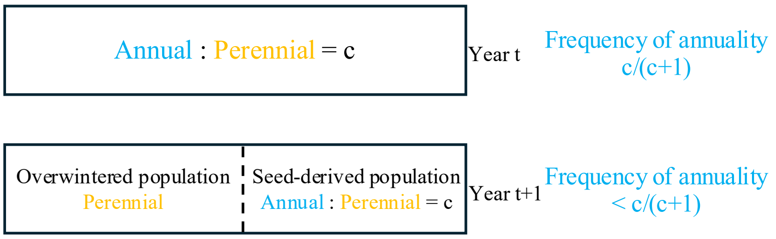


**Supplementary Figure 4: Frequency change of annuality vs. perenniality over time.** In a mixed population with both annuals and perennials, the ratio of the annual to perennial individuals should theoretically remain stable in the seed-derived cohorts. However, perennials possess the additional advantage of overwintering and contributing to the gene pool of the subsequent generations. This would predict a progressive increase of the perenniality allele frequency over time.

**Supplementary Table 1: Information for the samples collected from South Korea.**

| Site | Koseong Bay (KB) | | Jindong Bay (JB) | |
|---|---|---|---|---|
| Latitude | 34°56'N | | 35°7'N | |
| Longitude | 128°14'E | | 128°33'E | |
| Sub-population code | KB.S(hallow) | KB.D(eep) | JB.S(hallow) | JB.D(eep) |
| Sub-population type | Perennial | Annual | Perennial | Annual |
| Seagrass area (ha) | 0.38 | 1.48 | 1.21 | 0.80 |
| Water depth (m) | 0.8 | 4.5 | 2 | 4.5 |

**Supplementary Table 2: Information for the clonemates.**

| Genet | Population | Clonemates | |
|---|---|---|---|
| G01 | JB.S | MJPV1 | MJPV3 |
| G02 | JB.S | MJPV11 | MJPV15 |
| G03 | JB.S | MJPV17 | MJPV2 |
| G04 | KB.S | KSPV11 | KSPV16 |
| G05 | KB.S | KSPV2 | KSPV3 |
